# Global meta-analysis reveals urban-associated behavioral differences among wild populations

**DOI:** 10.1101/2024.05.17.594668

**Authors:** Tracy T. Burkhard, Ned A. Dochtermann, Anne Charmantier

## Abstract

Urbanization is causing fundamental changes to natural environments, effecting rapid and substantial adaptive phenotypic change in wild populations. While a large body of work has investigated how urbanization may shape *interspecific* variation in behavioral traits, such as via urban environmental filtering, no study has yet quantitatively assessed global patterns of urban-associated *intraspecific* behavioral variation. Here, we conducted a phylogenetic meta-analysis to assess urban-associated behavioral differences in wild populations of birds, mammals, amphibians, reptiles, and insects. We focused on four commonly measured behaviors (boldness, aggression, activity, and exploration) and extracted paired urban-nonurban effect size estimates for phenotypic means and variances (*k* = 278), behavioral repeatability (*k* = 13), and behavioral correlations (*k* = 14) from 80 studies. We found clear evidence that urban populations exhibit heightened average boldness, aggression, exploration, and activity compared to nonurban conspecifics, a result that was robust among species, geographic region, and ecological niche. Further, our results suggested that “generalist” species have the strongest behavioral responses. Conversely, we did not find strong evidence linking urbanization to changes in phenotypic variation, behavioral repeatability, or behavioral correlations. Our results summarize data from a rapidly evolving field in urban ecology and demonstrate geographically and taxonomically widespread differences in behavior between urban and nonurban populations, suggesting parallel natural selection across urban species and populations.

## Introduction

Human activity is rapidly and critically changing natural environments, leading to both unprecedented losses of species (Braje and Erlandson, 2013; Turvey and Crees, 2019; Waters et al., 2016) and alteration of communities (McKinney, 2008; Sol et al., 2014). Within species, urbanization and its associated conditions impose survival challenges and selective pressures on organisms (Otto, 2018). These different conditions frequently correspond to behavioral changes, which are likely necessary for survival in urban environments (Lowry et al., 2013; Miranda et al., 2013; Sol et al., 2013).

Behavioral changes are important for surviving in urbanized environments due to the influence of behavior on life history and fitness-related traits (Brehm and Mortelliti, 2023; Smith and Blumstein, 2008; Szabo and Ringler, 2023), such as the affinity for dispersal, reproductive investment, willingness to sample new resources, and more (Brehm and Mortelliti, 2023; Chapple et al., 2012; Cote et al., 2010; Fogarty et al., 2011; Moiron et al., 2020). Recent work in urban ecology has focused on behavioral differences between urban and nonurban conspecific populations in four behavioral traits—boldness, exploration, aggression, and activity—that can directly influence survival probabilities (Dall et al., 2012; Lowry et al., 2013; Réale et al., 2007; Sol et al., 2013). Unfortunately, whether patterns of urban-nonurban population differences in behaviors are consistent across taxa or geographic regions has not yet been tested. Similarly, whether total behavioral variation or behavioral repeatability (*i.e.,* the proportion of total phenotypic variance explained by among-individual variance, Stoffel et al., 2017), differs between urban and nonurban populations has not been tested across species. Finally, whether among-individual correlations (*i.e.,* behavioral syndromes, Dingemanse et al., 2012), are broken, maintained, or reinforced in urban environments across species is likewise unknown (Ouyang et al., 2018). These represent significant gaps in the literature because changes in behavioral means, variations, repeatability, and syndromes would provide important and complementary insights into both how behaviors are expressed in response to urbanization and their potential to evolve.

To date, no study has quantitatively compared the behavior of urban and non-urban populations across species around the world. However, some conceptual models, reviews, and related meta-analyses allow for general predictions. In a review of 29 empirical studies comparing urban and nonurban populations of wild birds, Miranda and colleagues found that 6 out of 9 studies reported increased aggression in urban populations and that 5 out of 6 studies reported increased boldness in urban populations; however, limited sample sizes prevented further statistical analysis of these differences (Miranda et al., 2013). Similarly, Ritzel and Gallo found that 7 out of 9 empirical studies reported increased aggression, exploration, and boldness in urban populations (Ritzel and Gallo, 2020). By contrast, in a recent meta-analysis focusing broadly on anthropogenic climate change, Gunn and colleagues found no evidence of clear directional changes in behaviors in relation to a composite category encompassing human disturbance, fishing pressure, and anthropogenic noise in addition to urbanization (Gunn et al., 2021).

In this study, we conducted a phylogenetic meta-analysis to estimate patterns of urban-associated differences in four behavioral traits: boldness, aggression, activity, and exploration. To help us draw meaningful comparisons across species and study designs, we re-interpreted behavioral data based on a common set of operational definitions (Réale et al., 2007) independent of interpretations and terminology originally made by study authors. Our main goal was to focus on paired urban-nonurban comparisons of conspecific terrestrial vertebrates and invertebrates to address the following specific questions: 1) Is urbanization associated with differences in average behavioral response? 2) Is urbanization associated with differences in behavioral variation, as measured by total phenotypic variation and repeatability? 3) Is urbanization associated with changes in the direction or strength of behavioral correlations? 4) Are behavioral responses to urbanization consistent across taxa, geographic region, and ecological niche?

Based on previous research (*e.g.,* Lowry et al., 2013; Miranda et al., 2013; Sol et al., 2013), we predicted that urban populations would exhibit heightened boldness, aggression, activity, and exploration. Further, we predicted that urban populations would display greater total behavioral variation relative to nonurban conspecific populations, potentially in response to relaxed selection, increased mutation rate, or developmental changes (*e.g.,* Capilla-Lasheras et al., 2022; Thompson et al., 2022). Likewise, we predicted that urban populations would have greater behavioral plasticity, as indicated by lower repeatability, if behavioral plasticity is adaptive in heterogeneous urban habitats, as hypothesized by several reviews (*e.g.,* Lowry et al., 2013; Sol et al., 2013). We also predicted that the strength of behavioral correlations would be less in urban populations relative to nonurban populations. Finally, we tested whether ecologically “generalist” species, which are hypothesized to be better equipped for urban environments (*e.g.,* Bonier et al., 2014; Ducatez et al., 2018; Lokatis et al., 2023; Sorace and Gustin, 2009) had different behavioral responses to urbanization than more specialized species.

## Methods

This study was initiated via a preregistered research plan (Open Science Framework; https://doi.org/10.17605/OSF.IO/GSXBM). We explain deviations from our preregistration in the supplementary materials. Briefly, our primary deviations were due to being unable to perform some pre-planned analyses given the available data. The final data composition also required analyses examining the potential for taxonomic bias in estimation.

### Systematic review and data extraction

We conducted literature searches following guidelines by the Preferred Reporting Items for Systematic Reviews and Meta-Analyses (PRISMA) (Page et al., 2021). Searches were completed prior to preregistration to roughly determine the types and amount of data available. We conducted four searches using the Web of Science Core Collection on September 8, 2022. We limited each search to the Science Citation Index Expanded database (SCI-EXPANDED, 1900 – present). Full details and search terms can be found in the supplementary materials.

The Web of Science (WoS) literature searches produced 1323 studies. We also identified 87 relevant studies based on work cited in three published reviews and meta-analysis. It is now known that different search engines (*e.g.,* Google Scholar, WoS) recover incomplete or inconsistent lists of articles due to institutional licensing, biases towards research from Western countries, and more (Gusenbauer, 2022; Pozsgai et al., 2021; Tennant, 2020). Thus, we also included 4 relevant papers identified by a colleague (MJ Thompson) that were not identified by WoS literature search. This resulted in 1086 unique studies (<u>Fig. S1</u>).

We excluded studies that did not meet our preregistered inclusion criteria (Supplemental materials). We also identified papers that analyzed the same or overlapping datasets, such as those from studies by the same lab or sets of authors that analyzed a longitudinally growing dataset. For these datasets (*N* = 8 studies), we included only the study using the dataset with the largest sample size, which was in all cases the most recently published study. If a study met the screening criteria, we extracted behavioral trait means, standard deviations (SD), and sample sizes (N) from the text, tables, or supplemental files. For cases where the standard error (SE), but not SD, was provided, we calculated standard deviation as 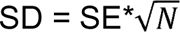, where *N* was the sample size. If these statistics were not available within either the article or supplemental files, we contacted the corresponding author(s) of the study for these estimates (*N* = 10). Authors of eight of these ten studies provided us with raw data or the necessary statistics and were subsequently included in analyses. When authors did not respond to our request (*N* = 2), we extracted mean and standard deviation or error estimates from available figures using a digital ruler (Van der Mierden et al., 2021). When available, we also extracted available repeatability coefficients (τ, unadjusted or adjusted for fixed effects, with preference for unadjusted, Nakagawa and Schielzeth, 2010) and phenotypic correlation coefficients (*r*).

In addition to extracting quantitative behavioral data, we recorded the following information for each study: common species name, species name, taxonomic class, geographic region, behavioral assay (*e.g.,* open field test, flight initiation distance), behavioral metrics (*e.g.,* distance run, quadrants explored, latency to flee), number of urban and nonurban populations, and whether behaviors were measured from wild animals tested or observed in the field, wild-caught animals tested in captivity, or from wild-derived animals bred and tested in a common environment. Behaviors were categorized based on operational definitions following Reale et al. (2007, Supplemental materials). Using fixed operational definitions reduced the possibility of naming fallacies affecting our results.

In our sample of studies, there was not a common metric for urbanization across studies. We instead relied on designations of “urban” and “nonurban” environments as provided by the authors and made the assumption that these urban-nonurban comparisons were relevant to the authors’ study systems.

### Collection of moderator data

We classified species as urban exploiters, adapters, or avoiders based on published designations in birds (Bonier et al., 2014; Croci et al., 2008) and mammals (Santini et al., 2019). To categorize each species’ ecological niche, we collected data on dietary type (*e.g.,* omnivore, carnivore, plant and seed, fruit and nectar, or invertebrate), dietary category (specialist or generalist), and temporal activity type (nocturnal, diurnal, or cathemeral) from open-access databases, using the EltonTraits 1.0 database (Wilman et al., 2014) for birds and mammals, the “amphibio” dataset from the traitdata R package for amphibians and reptiles (version 0.0.1, Meiri, 2019, 2018; Oliveira et al., 2017b, 2017a; RS-eco, 2022), and data from (Digweed, 1994; Luff, 1978; Stone et al., 2002; Tseng et al., 2018; Vucic-Pestic et al., 2010; Williams, 1959; Zygmunt et al., 2006) for invertebrates.

### Calculation of effect sizes

We calculated between-group standardized mean differences (SMD, (Hedges, 1981)) and differences in log standard deviation (Δln(SD), Senior et al., 2020) to investigate differences in mean behavioral traits and total behavioral variation, respectively. We calculated SMD as Hedges’ g corrected for bias and sample size (Hedges, 1981) using the metafor package (Viechtbauer, 2010). We calculated differences in ln(SD) as the unbiased estimator of the natural logarithm of the population standard deviation (ln a) following (Nakagawa et al., 2015):

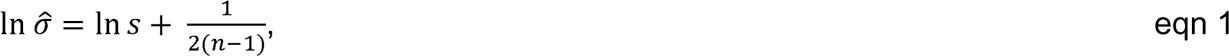

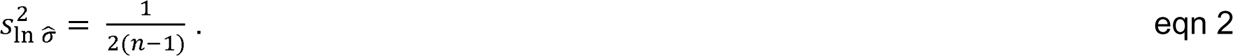

When available, we also recorded paired urban-nonurban estimates of repeatability (τ) and behavioral correlations (*r*, <u>Fig. S1</u>). Due to a small sample size of reported repeatability coefficients, we pooled unadjusted and adjusted repeatability coefficients in our analyses of repeatability. Likewise, we pooled between-individual correlation coefficients and total phenotypic correlations for analyses of behavioral correlations.

In total, we were able to calculate 285 effect sizes for both SMD and Δln(SD). Of these, we excluded 7 effect sizes because they had standard deviations equal to 0, ending with a final set of 278 effect sizes for both SMD and Δln(SD) (<u>Fig. S1</u>). We adjusted the sign of effect size estimates such that positive values indicated higher mean values *(e.g*., heightened boldness, exploration, activity, or aggression) in urban populations. In total, we changed the sign of 210 of 278 effect sizes. Repeatability and correlation coefficients were transformed for analyses using Fisher’s z transformation (Fisher, 1921) and back-transformed for interpretation.

### Data analysis

Meta-analyses of standardized mean differences (SMD), repeatability, and correlation coefficients were run using the metafor package, with visualizations produced using metafor and the orchaRd package (v2.0, Nakagawa et al., 2021). Each model was fitted assuming unstructured variance-covariance matrices. For analyses comparing Δln(SD), we used the MCMCglmm package (version 2.34, Hadfield, 2010). Full details for MCMCglmm models can be found in the Supplemental materials. For all models, we considered estimates to be statistically supported if they had a 95% confidence interval or, for Bayesian models, 95% credible intervals, that did not overlap zero. Phylogenetic trees to account for phylogenetic non-independence were built using phyloT (Letunic, 2015).

### Intercept-only models

We fitted phylogenetic meta-regression models with an intercept but no moderators to estimate the global effect size of urbanization for each component of behavioral differences (*i.e.,* SMD, Δln(SD), repeatability, and behavioral correlations, <u>Fig. S1</u>). We first fitted models that pooled behavior type for analyses of mean, behavioral variability, repeatability, and behavioral correlation combinations (*e.g.,* aggression-boldness, exploration-activity) for analyses of behavioral syndromes, thus estimating a global effect size for a general behavioral response, not for each behavioral type. We then fitted separate intercept-only models by behavioral type for analyses of mean, variance, and repeatability. For each intercept-only model, we included four random effects: study identity, observation identity, species, and phylogeny. Due to the limited and imbalanced sample sizes for each behavioral combination, we refrained from fitting behavioral-combination-specific models for syndromes.

Finally, to better understand how birds, which were the primary taxa in our data set (Fig. 1a), affected our results, we fitted separate intercept-only models for: 1) Aves only and 2) non-avian taxa for SMD and Δln(SD) datasets (<u>Fig. S1</u>).

**Figure 1.**
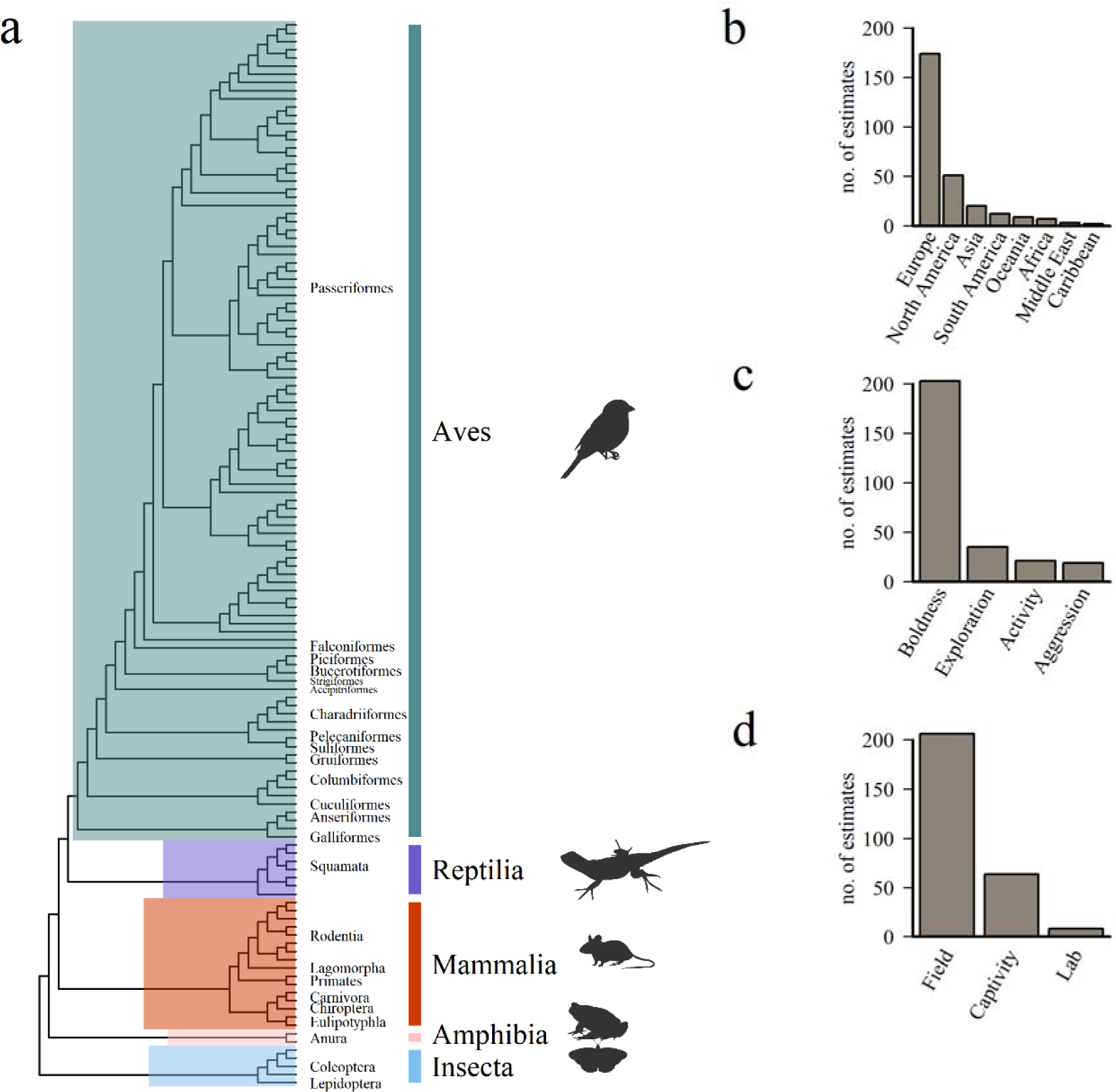
Taxonomic, behavioral, and geographic breadth of the meta-analysis. (a) Phylogenetic tree depicting all species represented in the 80 papers included in the meta-analysis. Branches are annotated with taxonomic orders represented in the study; the full list of species can be found in the Appendix. (b-d) Bar charts showing number of paired urban-nonurban observations for each category in the 80 papers included in the meta-analysis. (b) Our meta-analysis included a broad range of geographic regions but was biased towards studies of European populations. (c) Among the four behaviors studied, measures of boldness comprised the majority of observations. (d) Studies included estimates made on wild animals recorded in the field, wild-caught animals recorded in captivity, and lab-reared, wild-derived animals recorded in captivity.

### Moderator meta-regression models

For the SMD and Δln(SD) datasets, which had the largest sample sizes, we tested whether the relationship between urbanization and behavioral response differed among *a priori* selected moderators. We first fitted a “full” model that included each of the following moderators: region, provenance of data (whether the study populations were wild animals observed in the field, wild animals observed in captivity, or wild-derived animals bred and observed in captivity), urban tolerance, diet type, diet specialism, and temporal activity (<u>Fig. S1</u>). The full model was fitted on the largest available subset of observations with complete information for each of the moderators, comprising 162 effect sizes from 40 studies for SMD and for Δln(SD). As some moderator categories had few estimates, or categories had uneven sample sizes (*e.g.,* the dataset was composed entirely of observations of birds and mammals, Fig. 1a), we also tested each moderator in separate meta-regressions (Royauté et al., 2018; Vincze et al., 2017). All models had the same random effects as the global intercept-only meta-analyses.

### Heterogeneity

Heterogeneity estimates the consistency of the effect sizes, which can be inflated if studies greatly differ methodologically, taxonomically, or in geographic region (Higgins & Thompson, 2002; Higgins et al., 2003; Cooper & Hedges, 2009). For analyses done in metafor, we estimated heterogeneity (*I^2^*) using the *i2_ml* function in the orchaRd package (version 2.0, Nakagawa et al., 2021), following (Nakagawa and Santos, 2012; Senior et al., 2016). To calculate heterogeneity (*I^2^)* of Bayesian models, we divided the modal estimated variance by the total variance in the data (equations 24 and 25, Nakagawa and Santos, 2012). We considered *I^2^* values around 25%, 50%, and 75% as low, moderate, and high heterogeneity respectively, following convention set by (Higgins et al., 2003).

### Publication bias

Publication biases can occur when large effect sizes are overrepresented in the literature, *i.e.,* small-study bias, or when effect sizes diminish over time as published studies begin showing reduced support for a given hypothesis, *i.e.,* time-lag bias (Jennions and Møller, 2002; Nakagawa et al., 2022). We tested for publication biases in the full dataset (*N =* 80 studies) using several complementary methods. First, we visually inspected funnel plots (Sterne and Harbord, 2004) and ran mixed-effects meta-regression model tests for funnel plot asymmetry. We then fitted separate meta-regression models to test for evidence of small-study publication bias and time-lag publication bias. For small-study bias, we included the square-root of the inverse of the effective sample size as a moderator (Nakagawa and Santos, 2012). For time-lag bias models, we included the mean-centered year of study publication as a moderator (Jennions and Møller, 2002). We fitted these models for SMD, repeatability, and behavioral syndromes, and each model was fitted first on the complete data set (all 278 observations), then on observations restricted to birds (*k* = 200), and finally on observations excluding birds (*k* = 78).

## Results

### Summary of datasets

Our final datasets included 80 studies reporting a total of 278 effect sizes for standardized mean differences (SMD) and for log standard deviation differences (Δln(SD)), 13 studies reporting 29 paired urban-nonurban repeatability coefficients (τ), and 14 studies reporting 14 paired urban-nonurban correlation coefficients (*r*) (<u>Fig. S1</u>). The studies reporting repeatability and syndrome coefficients were a subset of the 80 studies we analyzed for SMD and Δln(SD).

Though the 80 studies included in our analyses spanned 130 different species, birds comprised the majority of observations (71.9%, <u>Fig. S1</u>, Fig. 1a). By contrast, ectothermic species collectively comprised fewer than 10% of total observations (amphibians, 1.4%; insects, 2.9%; reptiles, 4.3%). Our dataset was also strongly biased towards studies conducted in Europe and North America (Fig. 1b).

Behavioral trait types were unevenly represented across studies (Fig. 1c). For SMD and Δln(SD), boldness was the best studied behavioral trait (*k* = 203 effect sizes, 73.0% of total effect sizes), followed by exploration (35, 12.6%), activity (21, 7.6%), and aggression (19, 6.8%). This bias for boldness was directly caused by the taxonomic bias favoring birds, as 169 of the boldness effect sizes (83.2%) were made from observations in birds; of these, 160 (94.7%) were from measures of flight initiation distance (FID), which is a relatively easy field measure of boldness in birds. Excluding avian taxa, behavioral traits were more equally represented in our SMD and lnSD datasets: out of 78 total effect sizes, 34 were for boldness (43.6%), 21 were for exploration (26.9%), 20 were for activity (25.6%), and 3 were for aggression (3.8%). Most of the observations in our study came from wild animals measured in the field (*k* = 206) or wild-caught animals measured in captivity (*k* = 64), with few observations coming from wild-derived animals reared and tested in a common garden (*k* = 8, Fig. 1d).

Among studies that reported repeatability estimates, exploration and boldness were the best documented (exploration: 14 estimates, 48.3% of paired estimates; boldness: *k* = 10, 34.5% of paired estimates), followed by activity (3, 10.3%) and aggression (2, 6.9%). Of studies reporting repeatability, 14 reported unadjusted repeatability and 9 reported repeatability adjusted for fixed effects.

We recovered 14 paired urban-nonurban correlation coefficients from 8 studies, 3 of which were reported as between-individual correlation coefficients, and 11 of which were reported as phenotypic correlations. Of the behavioral combinations reported, we recovered 6 paired correlation coefficients for aggression and boldness, 4 for boldness and exploration, 3 for activity and boldness, and 1 for activity and exploration.

### Is urbanization associated with changes in average behaviors?

Urban populations were both more aggressive (SMD [95% confidence interval] = 0.65 [0.05, 1.24]) and bolder (1.34 [0.65, 2.03]) than nonurban populations. Urban populations also tended to be more active (0.40 [-0.24, 1.05]) and more explorative (0.28 [-0.28, 0.84]) than nonurban conspecifics, though confidence intervals for both overlapped zero. These behavior-specific differences were reflected in an overall analysis, in which the standardized mean difference was positive, indicating a greater mean for behaviors of animals in urban versus nonurban populations (SMD estimate [95% CI] = 1.00 [0.32, 1.68]; Fig. 2a; Table1).

**Figure 2.**
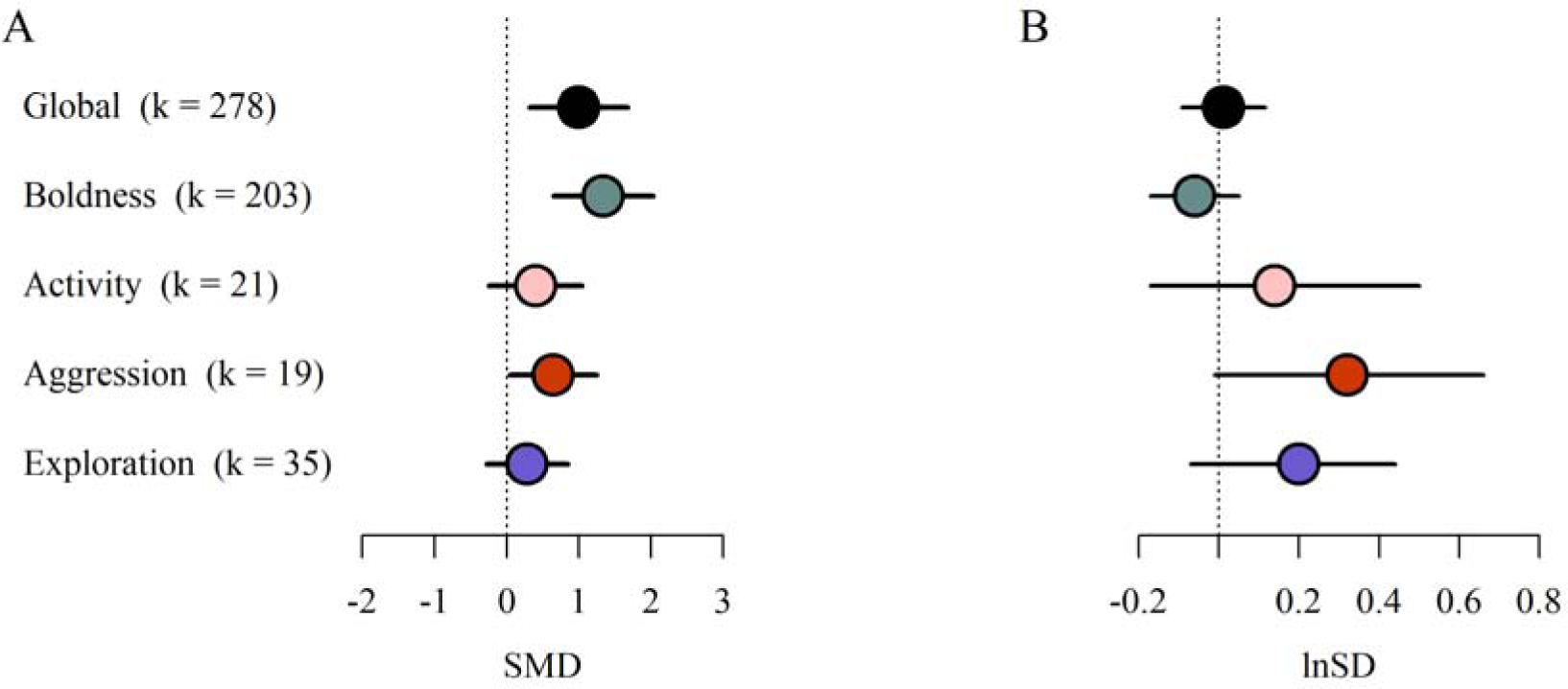
Forest plots of estimates for (a) standardized mean differences, SMD, and (b) differences in log standard deviation, Δln(SD). (a) Urban populations have greater mean behavioral responses than nonurban populations in all behaviors, but especially boldness and aggression. (b) Conversely, urban populations do not differ from nonurban populations in behavioral variation. Positive values on the x-axis represent higher mean (a) or variation (b) in urban populations than in nonurban populations. Black error bars indicate (a) 95% confidence intervals or (b) 95% credible intervals calculated from highest posterior density interval (HDPI); *k* provides the number of paired urban-nonurban estimates.

**Table 1.**
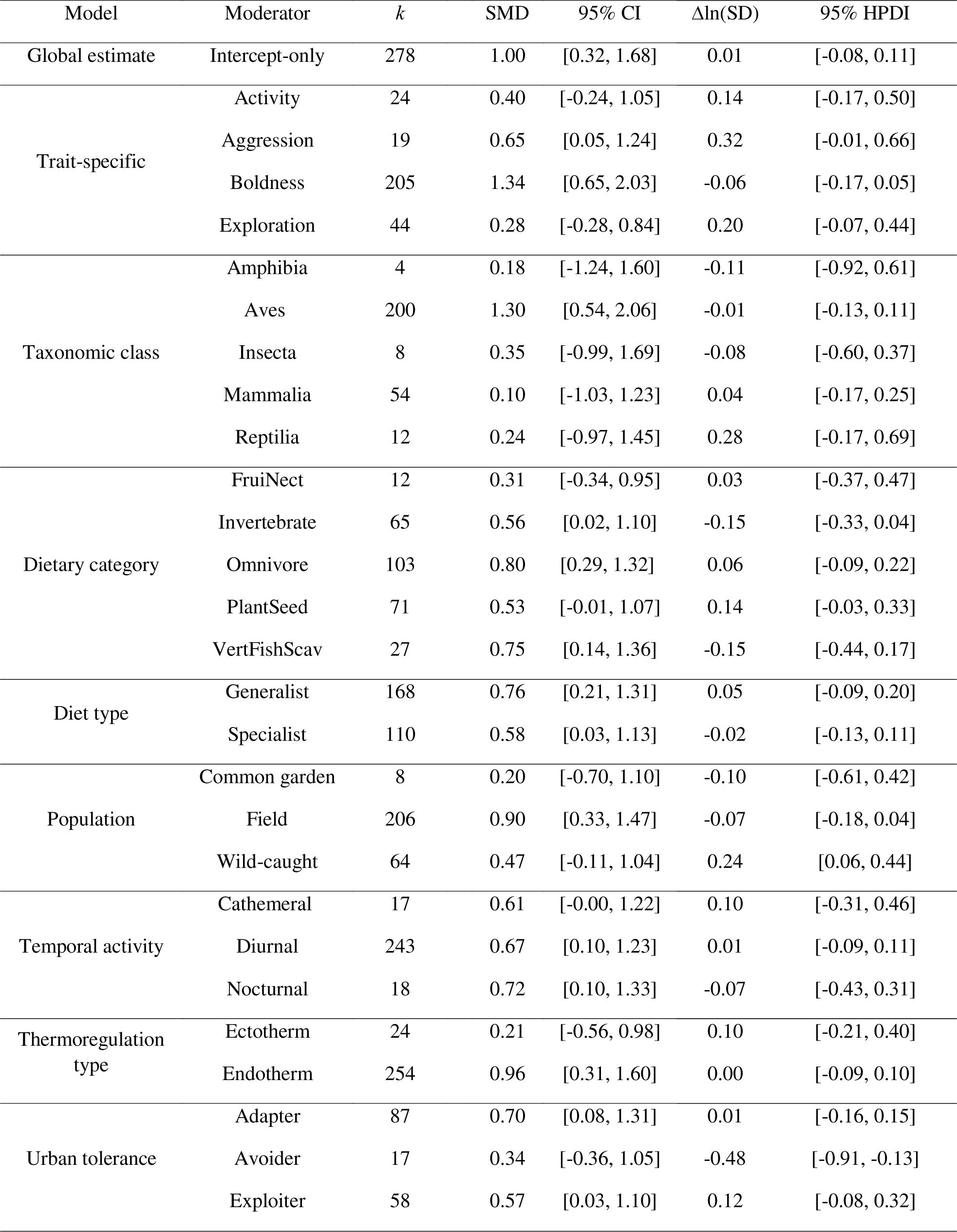
Estimates of SMD and Δln(SD) from our meta-analytic models. We show effect size estimates with associated 95% confidence intervals (SMD) or highest posterior density intervals (Δln(SD)) in brackets. *k* indicates the number of observations for each moderator.

Results for the avian subset of data likewise showed that urban populations of birds exhibited significantly greater averages for all four behaviors as compared to nonurban populations (though note that estimates for activity were based on a sample size of 1). Conversely, excluding birds from the analysis dampened the relationship between behavioral response and urbanization (SMD [95% CI] = 1.00 [0.32, 1.68] versus 0.51 [-0.04, 1.05] for non-avian species, <u>Table S1</u>). Only boldness remained significantly different between urban and nonurban non-avian populations (1.09 [0.44, 1.75]; <u>Table S1</u>).

### Is urbanization associated with changes to behavioral variation?

Urban populations tended to have more behavioral variation than nonurban populations in aggression (Δln(SD) [95% credible intervals]; 0.32 [-0.01, 0.66]), activity (0.14 [-0.17, 0.50]), and exploration (0.20 [-0.07, 0.44], Fig. 2b, Table 1). However, urban populations displayed less variation in boldness relative to nonurban populations (-0.06 [-0.17, 0.05], Fig. 2b, Table 1).

Importantly, Bayesian credible intervals for all behaviors overlapped zero, suggesting that these differences were not strong. We repeated these analyses in avian-only and non-avian-only subsets; differences in magnitude of phenotypic variation were not strong in either subset (<u>Table S2</u>).

We then focused on pairwise urban-nonurban comparisons of repeatability (Fig. 3, Table 2). Urban and nonurban populations had moderate and similar degrees of behavioral repeatability (back-transformed τ [95% confidence intervals]; nonurban τ: 0.36 [0.23, 0.47]; urban τ: 0.40 [0.28, 0.51]). Models parsing out specific behaviors revealed that the confidence intervals for urban and nonurban populations overlapped for each estimate, suggesting that there was no strong difference in repeatability between urban and nonurban populations for any behavior. Behaviors differed in their repeatability, with boldness having the highest repeatability for both urban and nonurban populations (back-transformed τ [95% CI]; urban τ: 0.49 [0.33, 0.63]; nonurban τ: 0.50 [0.33, 0.64]). Repeatability was also moderately high for activity (urban, 0.44 [0.18, 0.64]; nonurban, 0.35 [0.07, 0.57]) and exploration (urban, 0.35 [0.21, 0.49]; nonurban, 0.28 [0.12, 0.42]). Aggression had the lowest repeatability, and confidence intervals for both urban and nonurban populations overlapped zero (urban, 0.21 [-0.14, 0.51]; nonurban, 0.19 [-0.17, 0.51]).

**Figure 3.**
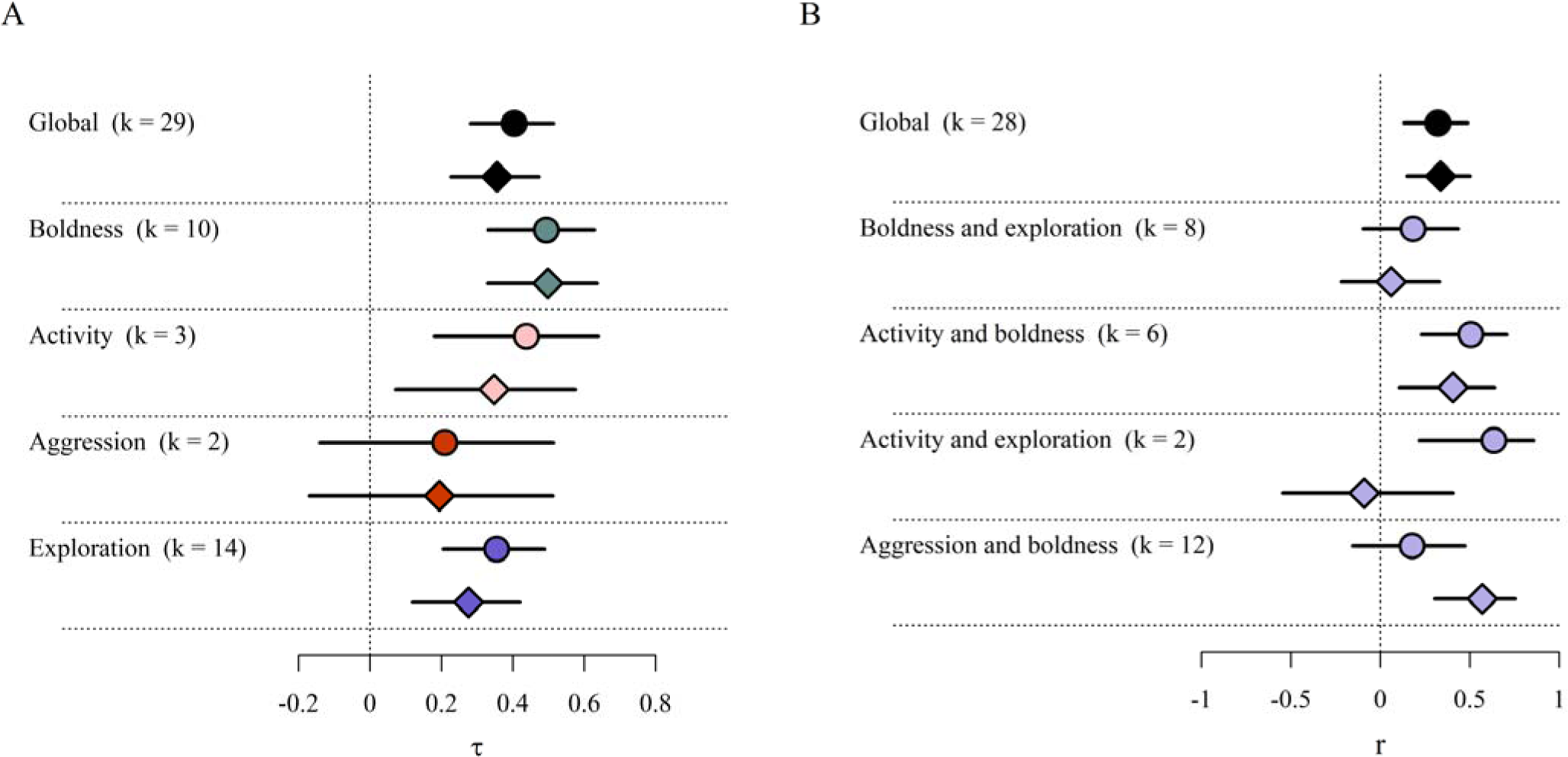
Forest plots of estimates for (a) repeatability correlation coefficients, ττ, and (B) behavioral syndromes, r. In both plots, circles = urban populations and diamonds = nonurban populations. Black error bars indicate 95% confidence intervals; k provides the number of observations.

**Table 2.** Estimates of repeatability with associated 95% confidence intervals. Estimates have been back-transformed. *k* indicates the number of observations for each moderator.

| Category | Population | *k* | $\tau$ | 95% CI |
| --- | --- | --- | --- | --- |
| Intercept-only | Urban | 29 | 0.4 | [0.23, 0.47] |
|  | Nonurban | 29 | 0.4 | [0.28, 0.51] |
| Activity | Urban | 3 | 0.4 | [0.18, 0.64] |
|  | Nonurban | 3 | 0.4 | [0.07, 0.57] |
| Aggression | Urban | 2 | 0.2 | [-0.14, 0.51] |
|  | Nonurban | 2 | 0.2 | [-0.17, 0.51] |
| Boldness | Urban | 10 | 0.5 | [0.33, 0.63] |
|  | Nonurban | 10 | 0.5 | [0.33, 0.64] |
| Exploration | Urban | 14 | 0.4 | [0.20, 0.49] |
|  | Nonurban | 14 | 0.3 | [0.12, 0.42] |

### Do nonurban and urban populations differ in behavioral correlations?

Behaviors were moderately and positively correlated in both urban and nonurban populations, and the strength of behavioral correlations did not differ between the two habitat types (back-transformed *r* [95% CI]; urban, 0.32 [0.13, 0.49]; nonurban, 0.34 [0.15, 0.50]; Table 3). Urban and nonurban populations varied in the strength of correlations with no consistency in which populations exhibited stronger correlations.

**Table 3.**
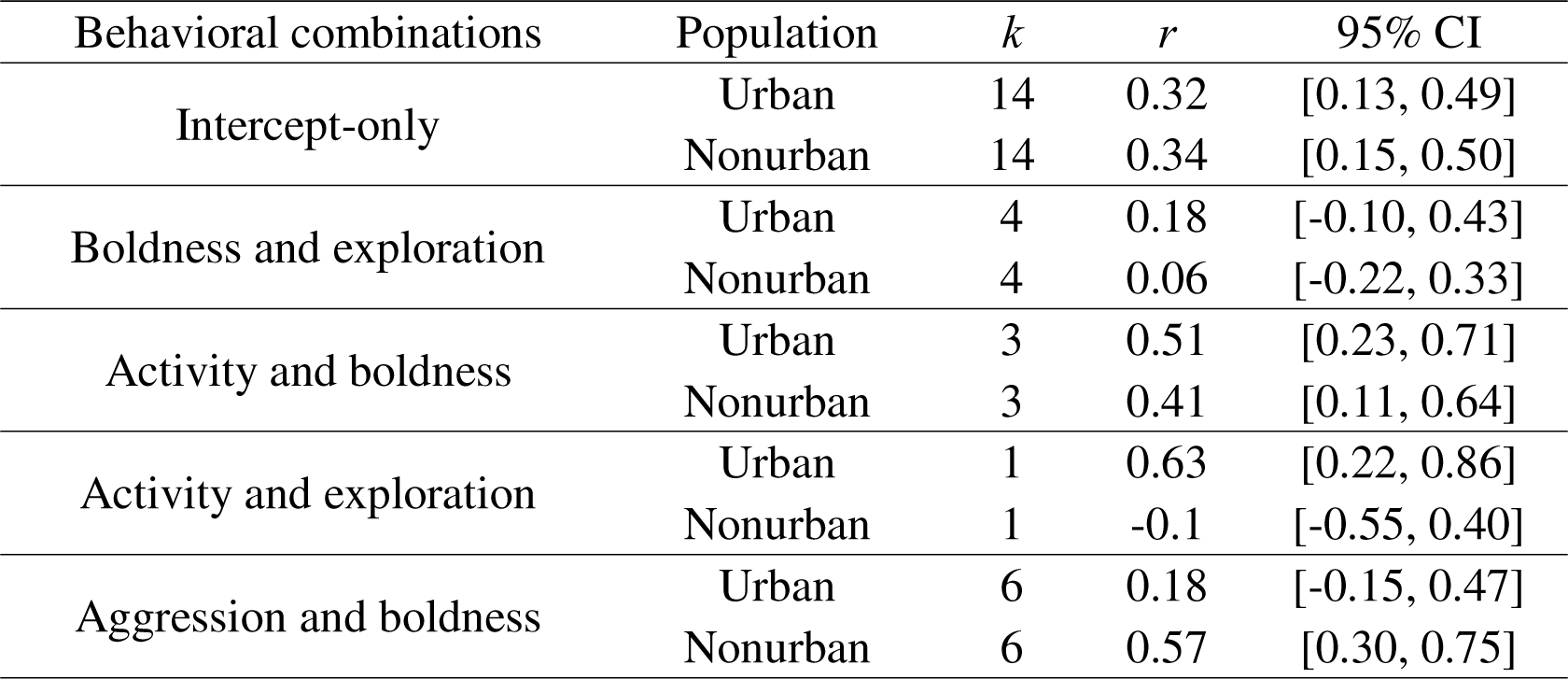
Estimates of behavioral correlations, *r,* with associated 95% confidence intervals. Estimates have been back-transformed; *k* indicates the number of observations for each moderator.

| Behavioral combinations | Population | *k* | *r* | 95% CI |
| --- | --- | --- | --- | --- |
| Intercept-only | Urban | 14 | 0.32 | [0.13, 0.49] |
|  | Nonurban | 14 | 0.34 | [0.15, 0.50] |
| Boldness and exploration | Urban | 4 | 0.18 | [-0.10, 0.43] |
|  | Nonurban | 4 | 0.06 | [-0.22, 0.33] |
| Activity and boldness | Urban | 3 | 0.51 | [0.23, 0.71] |
|  | Nonurban | 3 | 0.41 | [0.11, 0.64] |
| Activity and exploration | Urban | 1 | 0.63 | [0.22, 0.86] |
|  | Nonurban | 1 | -0.1 | [-0.55, 0.40] |
| Aggression and boldness | Urban | 6 | 0.18 | [-0.15, 0.47] |
|  | Nonurban | 6 | 0.57 | [0.30, 0.75] |

### Effects of moderators

For SMD, effect sizes for all moderators were positive. Specifically, urban populations exhibited increased average behaviors relative to nonurban populations regardless of class, ecological category, or urban tolerance category, though confidence intervals overlapped zero in many cases, particularly in categories with few samples (<u>Fig. S1</u>, Table 1). Interestingly, generalist species filling broader ecological niches—such as dietary omnivores and carnivores, endotherms, generalist dietary types, and urban adapters—tended to have the greatest urban behavioral differences in comparison with specialist species filling narrower niches (Fig. 4a).

**Figure 4.**
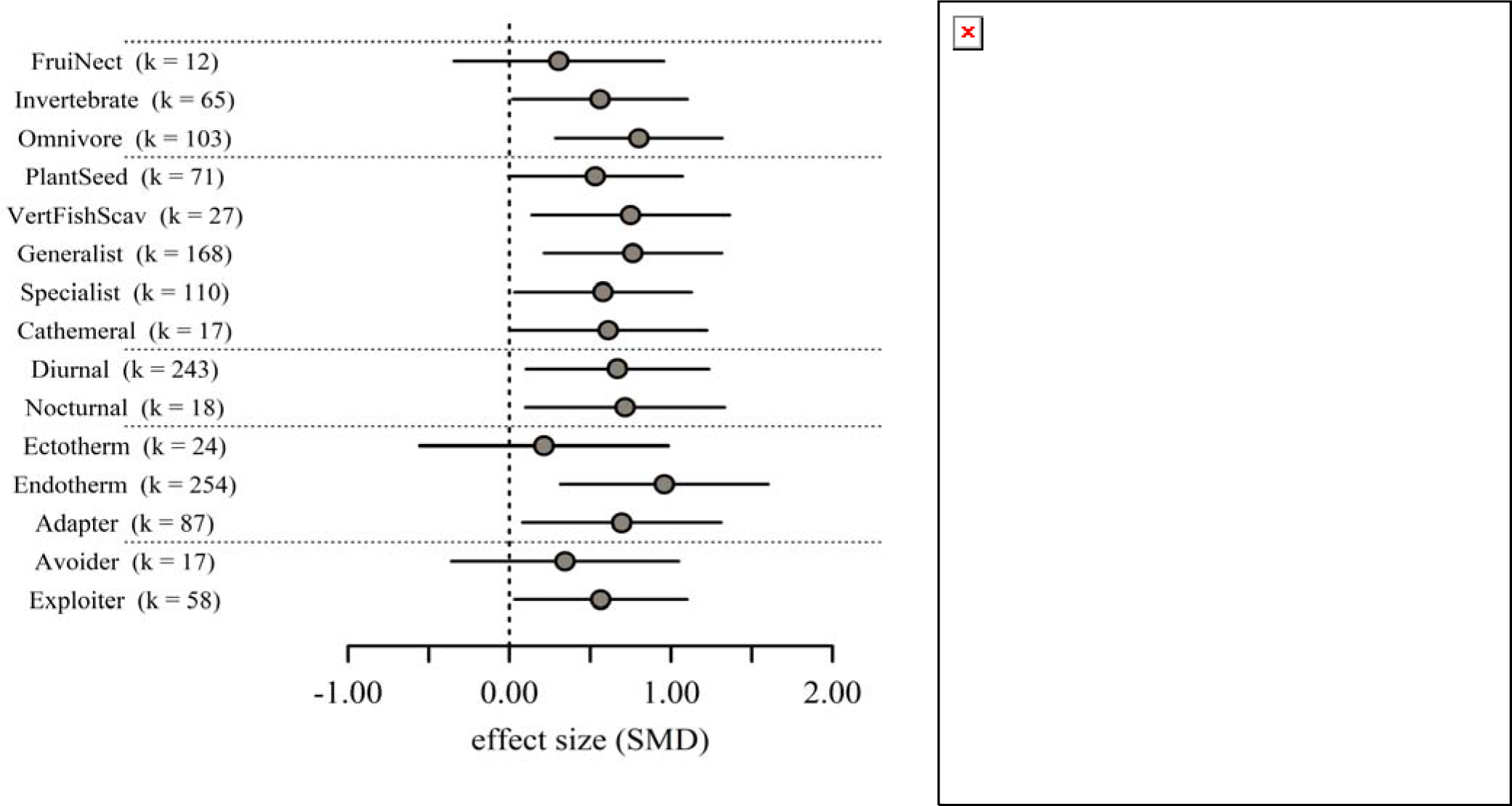
Moderator forest plots of estimates for (a) standardized mean differences, SMD, and (b) differences in log standard deviation, Δln(SD). Positive values on the x-axis represent higher mean (a) or variation (b) in urban populations than in nonurban populations. Black error bars indicate (a) 95% confidence intervals or (b) 95% credible intervals calculated from highest posterior density interval (HDPI); *k* provides the number of paired urban-nonurban estimates.

In contrast, we did not identify consistent patterns of differences in phenotypic variation, Δln(SD), among taxonomic classes or ecological categories (Fig. 4b). Among tolerance categories, however, we found evidence that “avoider” species exhibited significantly less phenotypic variation in urban populations relative to nonurban populations (Table 1).

### Heterogeneity

Total heterogeneity was high (>90%) for all models (Table S3). Study-level heterogeneity ranged from low to high, with studies reporting behavioral syndromes having the lowest study differences and studies reporting SMD having substantial differences explained by study differences (75%). Heterogeneity due to phylogenetic or species-level effects ranged from 13-18%, indicating a non-negligible but weak phylogenetic signal.

### Assessment for publication bias

We investigated the presence of small-study and time lag effects for all 80 studies included in our meta-analysis. Funnel plots were asymmetric, indicating publication bias, especially for SMD (Fig. S2). Meta-regression tests helped clarify these biases: while we found no evidence for time lag biases (*β* [95% CI]; *β*_Year_ = 0.31 [-0.69, 0.06]), we did find strong evidence for small study bias (*β*_N_ = -0.94 [-1.64, -0.24]), indicating that smaller studies may systematically overestimate the observed effects compared to larger studies.

## Discussion

In this study, we used phylogenetically controlled meta-analyses to examine urban-associated behavioral differences in wild animals, focusing on four behaviors: aggression, boldness, exploration, and activity. We compared urban and nonurban average behavioral traits, phenotypic variation, behavioral repeatability, and correlations. Our results revealed that urban populations were on average bolder, more aggressive, more explorative, and more active relative to nonurban conspecifics, with boldness showing the most robust response. This pattern held across species from different ecological categories, with all categories showing urban-associated increases in average behavioral traits. Further, our results suggested that generalist species had the greatest behavioral responses to urbanization. Conversely, we found no evidence that urbanization was clearly associated with differences in behavioral variation, repeatability, or correlation strength, although there were some differences in correlations among trait pairs. Our findings provide evidence that urbanization has a taxonomically and geographically widespread effect on behavioral phenotypes.

### Evidence for increased average behaviors

Urbanization has been extensively linked to shifts in mean phenotypes, including shifts in morphology (*e.g.,* limb structure in Anolis lizards, Winchell et al., 2023; melanism in peppered moths, Kettlewell, 1958, 1956), life history strategies (*e.g.,* clutch size and lay date in birds, Capilla-Lasheras et al., 2022; Sepp et al., 2018), and physiology (*e.g.,* thermal tolerance in ants, Diamond et al., 2017; stress response in juncos, Atwell et al., 2012). Here, we found compelling quantitative evidence that urbanization is also linked to increases in mean behavioral traits (*i.e.,* heightened boldness, exploration, activity, and aggression), thereby expanding our understanding of urban-associated phenotypic shifts and corroborating qualitative findings from previous reviews (Lowry et al., 2013; Putman and Tippie, 2020; Ritzel and Gallo, 2020; Sol et al., 2013). Urban populations showed increases in all four behavioral traits, particularly boldness and aggression, but the magnitude of this response varied across taxa. When we considered only avian taxa, which comprised 71.9% of effect sizes, all four behavioral traits showed strong increases in urban populations (note, however, that results for activity were based only on one paired urban-nonurban effect size, <u>Table S1</u>). Conversely, directional urban-specific behavioral differences were less evident among non-avian species: boldness was the sole behavior expressing a significant increase in urban populations, and no other behaviors differed between urban and nonurban populations. While the lack of significant behavioral change could indicate a true absence of urban behavioral change in non-avian species, *i.e.,* a biological non-result, an alternative explanation is that a limited sample size reduced our ability to detect nonurban-urban differences in non-avian species. Indeed, though we strove to extend previous work by synthesizing estimates from mammals, amphibians, reptiles, and invertebrates, these non-avian species were still poorly represented in our final dataset (Fig. 1a), reflecting ongoing taxonomic representation issues widely observed in animal behavior and biodiversity research (Rosenthal et al., 2017; Troudet et al., 2017).

Boldness was substantially higher in urban populations, a robust pattern observed across both non-avian and avian taxa. Several factors could contribute to this phenomenon. First, being bolder and taking more risks may increase an individual’s likelihood of settling in urban areas, either because bold individuals have a greater propensity for dispersal, either actively (Bégué et al., 2023; Dingemanse et al., 2003; Fraser et al., 2001) or passively (*e.g.,* “uptake”, Chapple et al., 2012), or because bold animals prefer specific habitat characteristics that may be present in an urban environment (*e.g.,* personality-matching habitat choice, Holtmann et al., 2017; Schirmer et al., 2019). Alternatively, though not mutually exclusively, increased boldness may be an adaptive plastic or genetic response in urban environments. Increased boldness may allow urban individuals to be less sensitive to human presence (Uchida et al., 2019) or give them a competitive edge in environments with limited or low-quality resources (Clermont et al., 2023; Tremmel and Müller, 2013) or changed predator communities (Ward-Fear et al., 2018). Additional studies are clearly needed to help disentangle the drivers of urban-associated boldness, and future work incorporating migration-dispersal dynamics and assessment of fitness benefits of behavioral responses will yield valuable insights on the causes and consequences of increased boldness in urban populations.

### No evidence for differences in behavioral variation

We did not find evidence suggesting that urban and nonurban populations differ in behavioral variation (Δln(SD), Table 1). This result contrasts with findings from two recent meta-analyses reporting urban-associated increased phenotypic variation in a variety of life history and morphological traits populations (Capilla-Lasheras et al., 2022; Thompson et al., 2022) but is consistent with large-scale studies demonstrating a lack of clear patterns. Specifically, in a large-scale analysis of phenotypic variation spanning 4507 effect sizes, 196 studies, and 177 species, Sanderson and colleagues found that phenotypic variation did not differ in a generalizable way by either trait type or human disturbance (Sanderson et al., 2023). Despite the absence of general patterns, it is still valuable to explore how phenotypic variation responds to urbanization within specific cases (*e.g.*, species, phenotype, urban characteristic) to better understand the evo-ecological dynamics underlying urban-associated phenotypic change.

### Implications for behavioral plasticity and evolutionary constraints

Urban and nonurban populations did not differ in repeatability, *i.e.,* the amount of phenotypic variance attributable to among-individual variance. Instead, we found evidence that in both urban and nonurban environments, the repeatability of each behavior ranged from low (τ *=* 0.19) to moderately high (τ = 0.50), with boldness being the most repeatable behavior. These values fall within the ranges of previously reported repeatabilities for “personality”-related behaviors (average τ, range; Dochtermann et al., 2014: 0.62 (0.37 – 0.83); Bell et al., 2009: 0.37 (-0.22 – 0.97). One notable difference in our study, however, is that aggression fell on the lower side of this range for both urban and nonurban populations (urban τ = 0.21, nonurban τ = 0.19), in contrast to previous reports of high repeatability for aggression (Dochtermann et al., 2014, τ = 0.65; Bell et al., 2009: τ = 0.48). This contradictory finding could be explained by the fact that in a focal individual, aggressive behavior is highly dependent on the social context (in particular, an opponent) and often influenced by indirect genetic effects (*e.g*., Bailey et al., 2018). Alternatively, since we independently assigned behaviors to categories, prior meta-analyses and individual studies may have conflated multiple different behaviors under the aggression label. For example, aggression is sometimes defined as agonistic interactions with conspecifics (Oyegbile and Marler, 2005), heterospecifics (Vullioud et al., 2013), or humans (Class et al., 2014), and these behaviors are not necessarily be equivalent (Gupta et al., 2019; Peiman and Robinson, 2010).

Given that repeatabilities provide an upper limit for heritability, our results imply that both urban and nonurban populations likely retain heritable variation on which selection can act. This is a very tentative conclusion, however, for three reasons: (1) there were very few studies that reported repeatability; (2) differences in repeatability are difficult to interpret given that they may differ due to divergence in both among- and within-individual variances (Dochtermann and Royauté, 2019; Royauté and Dochtermann, 2021; Wilson, 2018); and (3) repeatability is not informative as to a lower limit of heritability. Consequently, future research should prioritize the reporting of the variance components that contribute to repeatabilties and prioritize the estimation of additive genetic variation. For similar reasons, the potential for behavioral correlations to constrain evolutionary responses to differing urban and nonurban selective regimes remains unclear.

In summary, our results show that urbanization is associated with substantial changes to behavior. Our results also highlight that these changes are pervasive across geographic regions, ecological niche, and species. Future research determining whether these differences represent adaptations to urban environments will help clarify how selection has changed in the Anthropocene and to what degree organisms can respond to these changes.

## Acknowledgements

The authors thank PB D’Amelio and SK Smith for helpful comments on the manuscript and MJ Thompson for discussion of relevant literature. Further, we thank K Uchida, MJ Thompson, S von Merten, D Blumstein, J Baxter-Gilbert, S Breck, J Hyman, and M Hasegawa for kindly providing data for the analysis.

## Notes

### Competing Interest Statement

The authors have declared no competing interest.

