## Supplementary materials for "Global meta-analysis reveals urban-associated behavioral differences among wild populations"

### Methods

#### Deviations from preregistration

Our final meta-analysis deviated from the preregistration in several ways:

- First, we could not specify certain analyses beforehand due to the uncertainty regarding the available data and its scope.
- For studies comparing multiple populations along an urban gradient, we included estimates only for the most urban and least urban populations to maintain consistency.
- We were also unable to use degree of urbanization as a moderator because studies differed substantially in both the method used to assess urbanization and in whether they reported this method or measurement.
- We were unable to fit behavioral correlation-specific models for syndromes due to the insufficient number of studies.
- The limited number of studies on non-avian taxa restricted our ability to conduct separate analyses for each taxonomic class.

#### Literature search and criteria for inclusion

We conducted literature searches following guidelines by the Preferred Reporting Items for Systematic Reviews and Meta-Analyses (PRISMA) (Page et al., 2021). Each search was completed prior to preregistration to determine the types and amount of data available. We conducted four searches using the Web of Science Core Collection on September 8, 2022, limiting all searches to the Science Citation Index Expanded database (SCI-EXPANDED, 1900 – present) to improve the relevance of returned articles. We performed the following searches: (**1**) Topic = [(*list of taxa*) AND (*list of urban habitats*) AND (*list of nonurban habitats*) AND (*list of behaviors*) AND (animal wild)]; (**2**) Topic = [(*list of taxa*) AND (*list of urban habitats*) AND (*list of nonurban habitats*) AND (*list of behaviors*)] AND Research Areas = (Zoology, Behavioral Sciences, Ornithology, Evolutionary Biology, Entomology); (**3**) Topic = [(*list of taxa*) AND (*list of urban habitats*) AND (*list of nonurban habitats*) AND (*list of tests*)]; (**4**) Topic = [(*list of taxa*) AND (*list of urban habitats*) AND (*list of nonurban habitats*) AND (behavioral syndrome OR behavioural syndrome OR Bold* OR Explorat* OR Neophilia OR Neophobia OR Novel object)]; (**5**) Topic = [(*list of taxa*) AND (*list of urban habitats*) AND (*list of nonurban habitats*) AND (proactive NEAR/1 reactive) syndrome OR personality traits OR (pace of life NEAR/1 behavior)].

Each “*list*” search term contained a list of the keywords used to specify taxa, environment, and behavior or behavioral assay:

*list of taxa =* (mammal* OR wild OR animal OR bird OR vertebrate OR avian OR reptil* OR herp* OR amphibian OR anuran OR frog OR lizard OR insect OR arthropod OR spider OR snake OR invertebrate)

*list of urban habitats =* (anthropogenic disturbance OR human disturbance OR urban* OR city* OR town* OR metro*)

*list of nonurban habitats =* (rural OR farm* OR agricult* OR country OR countryside OR natural environment OR forest*)

*list of behaviors* = (affiliative OR affiliat* OR aggress* OR alarm OR anti-predator OR behavioural syndrome OR behavioral syndrome OR bold* OR defense OR escape OR explor* OR fear OR FID OR flight initiation distance OR gregarious* OR neophilia OR neophobia OR personality OR risk-taking OR shy OR sociability OR temperament OR vigilance)

*list of tests* = (open field OR novel object OR handling bag test OR hole board test OR (light NEAR/1 dark) test OR light-dark test OR predator presentation test OR predator stimulus test OR novel environment OR mirror image OR mirror test OR attack latency OR dyadic encounter OR diadic encounter)

These searches turned up 119, 835, 318, and 51 articles, respectively. We de-duplicated this list using the ‘Remove Duplicates’ function in Excel (Microsoft Office), followed by visual inspection of the author lists and titles. De-duplication resulted in a list of 1086 unique studies.

We read the title and abstract of each study in our de-duplicated list to screen for eligibility in our meta-analysis. Studies were included only if they recorded behavioral data for at least one urban population and one non-urban population and provided species-specific behavioral data (e*.g.,* not including ‘mixed flock’ or species lumped together). While our study focused on wild populations, not domesticated animals or laboratory strains, we included common garden studies in which founding members were caught from wild urban and non-urban populations. To prevent pseudoreplication, we identified papers that analyzed the same or overlapping datasets, such as those from studies by the same lab or set of authors. For these datasets (*N* = 8 studies), we included only the study using the dataset with the largest sample size, which was typically the most recently published study.

#### Defining behavioral categories

We initially focused on the behavioral traits of boldness, aggression, activity, exploration, and sociability because of the rich literature establishing their role on survival and reproductive success in a wide range of species (Koolhaas et al., 1999; Réale et al., 2010; Smith and Blumstein, 2008). Our literature searches returned no articles with observations of sociality in an urban context, so we omitted sociality from our analyses. We defined the four behaviors following language proposed by (Réale et al., 2007): we defined “boldness” as individual reactions to risk, “exploration” as reactions to novel situations in the absence of risk, “activity” as the amount of movement of an individual, and “aggressiveness” as agonistic reactions by individuals to conspecifics. Note that under our study’s operational definitions, aggressive responses towards predators or humans are interpreted as measures of boldness. Finally, we recorded the behavioral methodology and metric(s) used in each behavioral observation (Supplemental Data Files).

#### Quantifying urbanization

Urbanization can be assessed by a multitude of factors, including human population density, infrastructure development, resource availability, pollution levels, and land use patterns (Melliger et al., 2018; Moll et al., 2020, 2019). In our literature review, we were unable to find a metric for urbanization that was used consistently across studies. For example, out of the 80 studies that comprised our final standardized mean differences (SMD) dataset, 20 defined urbanization as percent impervious cover, 8 defined urbanization as human population per km^2^, and 3 defined urban and nonurban populations as relative differences in total human population density. Rather than limiting our dataset to only those studies using a certain definition for urbanization, we instead relied on designations of “urban” and “nonurban” environments as provided by the authors and made the assumption that these urban-nonurban comparisons are relevant to the authors’ study systems.

#### Collection of moderator data

We classified species as urban exploiters, adapters, or avoiders based on published designations in birds (Bonier et al., 2014; Croci et al., 2008) and mammals (Santini et al., 2019). Urban exploiters rely on human-provided resources, adapters can live in and tolerate modified urban environments, and avoiders are unable to tolerate or live in urban environments (Blair, 1996; McKinney, 2002). In practice, these categories are often assigned by presence/absence data (Rodriguez et al., 2021; Santini et al., 2019). There are limitations to these categories, particularly with regards to the challenge of defining "urban" environments and the potential mislabeling of species (Fischer et al., 2015); further, we note that by our own criteria, all species included in our study were present in both urban and nonurban habitats. Nonetheless, we find these categories useful as they provide a simple framework for understanding how behavioral responses may differ among taxa with different tolerances of urbanization.

To categorize each species’ ecological niche, we collected data on dietary type, dietary category, and temporal activity type from open-access databases, using the EltonTraits 1.0 database (Wilman et al., 2014) for birds and mammals, the “amphibio” dataset from the traitdata R package for amphibians and reptiles (version 0.0.1, Meiri, 2019, 2018; Oliveira et al., 2017b, 2017a; RS-eco, 2022), and data from (Digweed, 1994; Luff, 1978; Stone et al., 2002; Tseng et al., 2018; Vucic-Pestic et al., 2010; Williams, 1959; Zygmunt et al., 2006) for invertebrates.

#### Data analysis

Meta-analyses of standardized mean differences (SMD), repeatability, and correlation coefficients were run using the metafor package, with visualizations produced using metafor and the orchaRd package (v2.0, Nakagawa et al., 2021). Each model was fitted assuming unstructured variance-covariance matrices.

For analyses comparing ∆ln(SD), we used the MCMCglmm package (version 2.34, Hadfield, 2010). We ran each model for 8,000,000 iterations with burn-in of 6,000,000 and selected a thinning interval of 2000 to reduce autocorrelation. This resulted in a posterior distribution sample size of 1000. For each model, we visually checked chains for convergence and mixing. We provided each model with an inverse gamma prior for random effects (V = 1, nu = 0.002) and used the default non-informative (flat) prior for fixed effects. We included population as a fixed effect, then calculated the difference between the urban and nonurban posterior distributions using the *describe_posterior* function from the bayestestR package (version 0.13.0, Makowski et al., 2019). To account for the mean-variance relationship, we also included the log mean as a fixed effect (Nakagawa et al., 2015). Inference on differences were based on posterior estimates and credible intervals (*i.e.*, highest posterior density intervals) overlapping zero or not. We then assessed urban effects per behavior and accounted for behavior-specific means and variances and possible correlations among behaviors by fitting a multivariate meta-analytic model with an unstructured variance-covariance matrix.

For all models, we considered estimates to be statistically supported if they had a 95% confidence interval or, for Bayesian models, 95% credible intervals, that did not overlap zero. Phylogenetic trees to account for phylogenetic non-independence were built using phyloT (Letunic, 2015).

### Supplemental figures

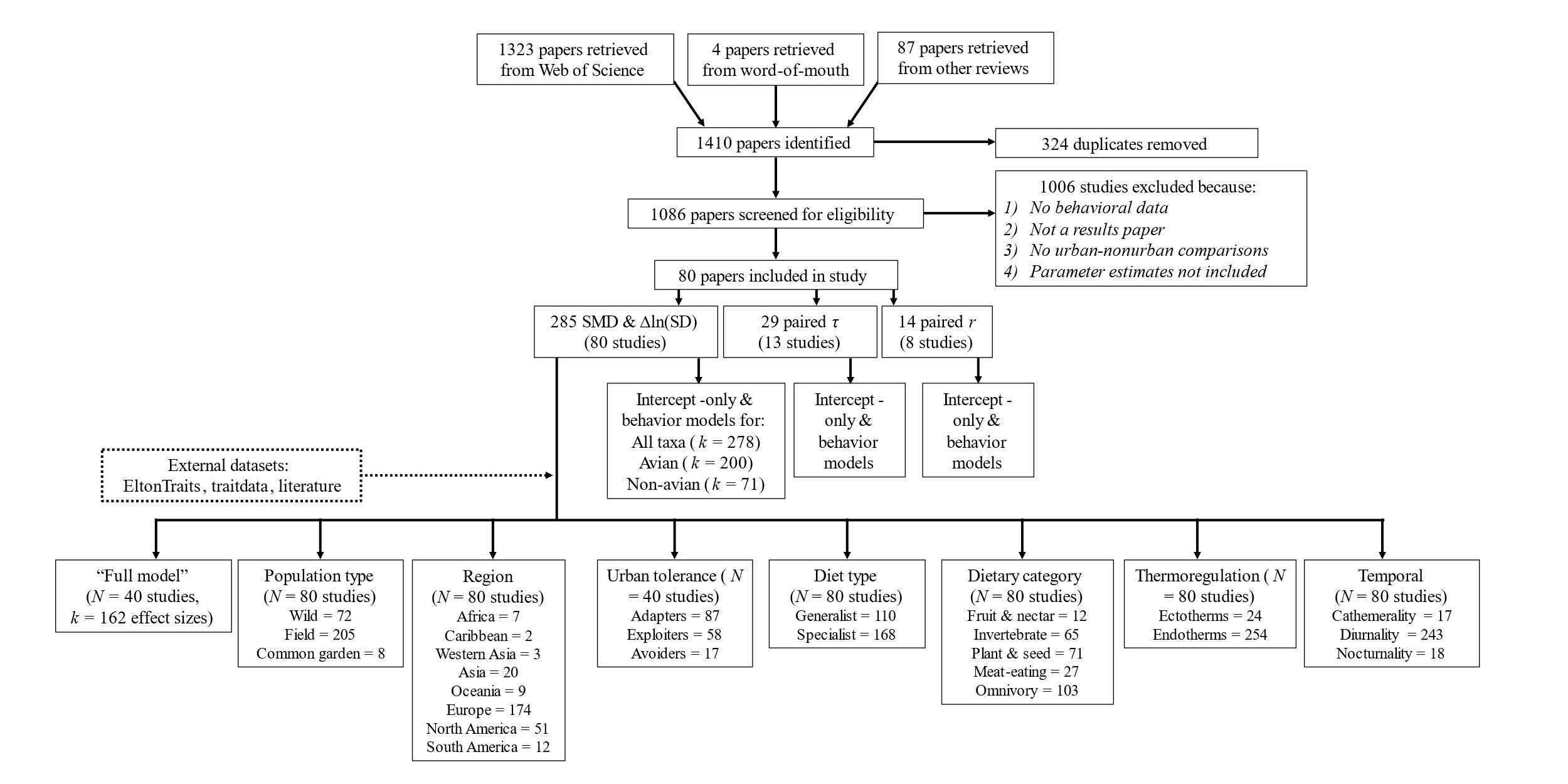

#### **Figure S1**

Prisma diagram

#### **Figure S2**

Funnel plots used to estimate publication bias in the full datasets for (A) SMD, (B) repeatability, and (C) behavioral syndromes. Meta-analytic means are indicated with dashed lines.

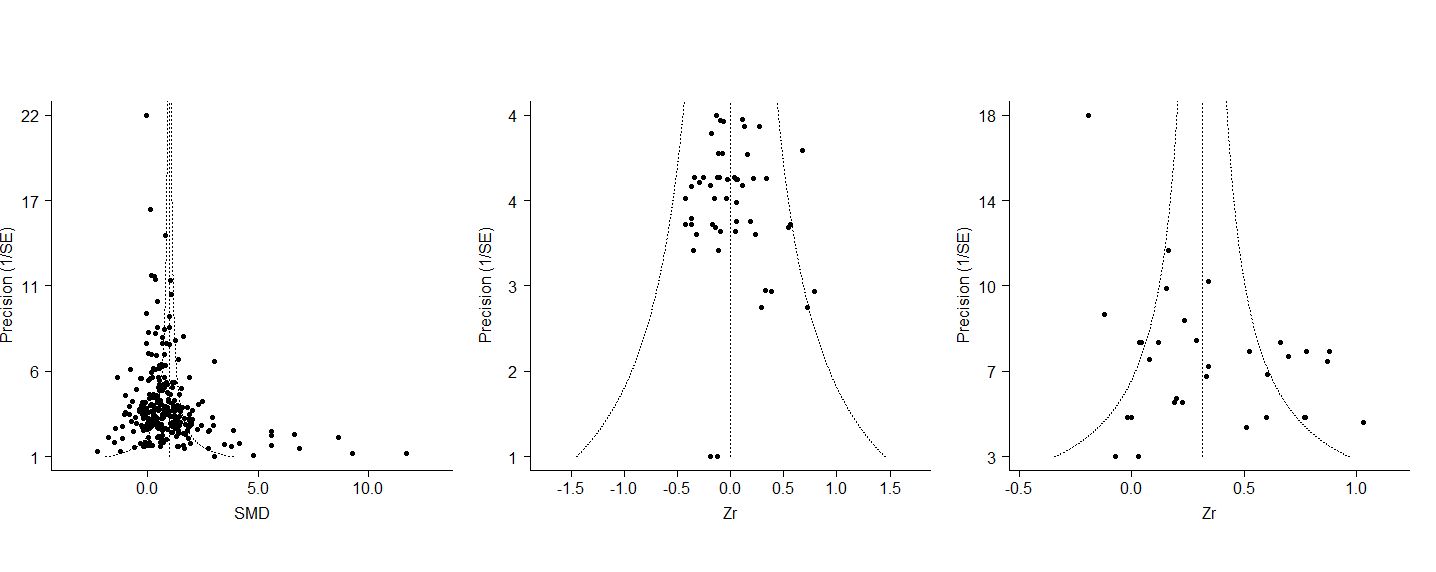

### Supplementary tables

#### **Table S1**

Standardized mean difference estimates for: 1) all taxa, 2) avian taxa, 3) non-avian taxa, including associated 95% confidence intervals (CI).

|  |  | Estimate | SE | Z | P | CI lower | CI upper |
| --- | --- | --- | --- | --- | --- | --- | --- |
| 1. All taxa | Global | 1.00 | 0.35 | 2.87 | 0.00 | 0.32 | 1.68 |
|  | Boldness | 1.23 | 0.35 | 3.58 | 0.00 | 0.56 | 1.91 |
|  | Activity | 0.93 | 0.38 | 2.45 | 0.01 | 0.19 | 1.68 |
|  | Aggression | 0.70 | 0.41 | 1.71 | 0.09 | -0.10 | 1.50 |
|  | Exploration | 0.56 | 0.37 | 1.53 | 0.13 | -0.16 | 1.29 |
| 2. Avian species | Global | 1.59 | 0.34 | 4.71 | <.0001 | 0.93 | 2.25 |
|  | Boldness | 1.78 | 0.39 | 4.62 | <.0001 | 1.03 | 2.54 |
|  | Activity | 1.54 | 0.73 | 2.12 | 0.03 | 0.11 | 2.96 |
|  | Aggression | 1.32 | 0.31 | 4.25 | <.0001 | 0.71 | 1.93 |
|  | Exploration | 0.94 | 0.30 | 3.11 | 0.00 | 0.35 | 1.53 |
| 3. Non-avian species | Global | 0.51 | 0.28 | 1.83 | 0.07 | -0.04 | 1.05 |
|  | Boldness | 1.09 | 0.33 | 3.27 | 0.00 | 0.44 | 1.75 |
|  | Activity | -0.26 | 0.19 | -1.39 | 0.16 | -0.63 | 0.11 |
|  | Aggression | 0.07 | 0.11 | 0.63 | 0.53 | -0.14 | 0.28 |
|  | Exploration | -0.11 | 0.16 | -0.66 | 0.51 | -0.42 | 0.21 |

#### **Table S2**

∆ln(SD) estimates for: 1) all taxa, 2) avian taxa, 3) non-avian taxa. 95% credible intervals included.

|  |  | Estimate | CI lower | CI upper |
| --- | --- | --- | --- | --- |
| All taxa | Global | 0.01 | -0.08 | 0.11 |
|  | Boldness | -0.06 | -0.17 | 0.05 |
|  | Activity | 0.14 | -0.17 | 0.50 |
|  | Aggression | 0.32 | -0.01 | 0.66 |
|  | Exploration | 0.20 | -0.07 | 0.44 |
| Avian species | Global | -0.08 | -0.22 | -0.08 |
|  | Boldness | -0.14 | -0.32 | 0.01 |
|  | Activity | -0.02 | -1.95 | 2.00 |
|  | Aggression | 0.44 | -0.07 | 0.94 |
|  | Exploration | 0.38 | -0.17 | 0.94 |
| Non-avian species | Global | 0.07 | -0.24 | 0.39 |
|  | Boldness | 0.02 | -0.38 | 0.46 |
|  | Activity | 0.17 | -0.36 | 0.73 |
|  | Aggression | -0.31 | -1.81 | 1.10 |
|  | Exploration | 0.08 | -0.46 | 0.65 |

#### **Table S3**

Heterogeneity

| Effect size | Descriptor | I2 Total | I2 study id | I2 obs id | I2 scientific name phylo | I2 scientific name |
| --- | --- | --- | --- | --- | --- | --- |
| SMD | Global, intercept-only | 98.41 | 75.32 | 9.36 | 13.73 | 0.00 |
| Syndrome | Global, intercept-only | 72.62 | 12.50 | 45.97 | 0.00 | 14.15 |
| Repeatability | Global, intercept-only | 76.02 | 22.39 | 34.97 | 2.05 | 16.60 |
| lnSD | Global, intercept-only | 99.73 | 78.83 | 20.82 | 0.003 | 0.004 |
